# Predicting the Degree of Distracted Driving Based on fNIRS Functional Connectivity: A Pilot Study

**DOI:** 10.1101/2022.01.25.477783

**Authors:** Takahiko Ogihara, Kensuke Tanioka, Tomoyuki Hiroyasu, Satoru Hiwa

## Abstract

Distracted driving is one of the main causes of traffic accidents. By predicting the attentional state of drivers, it is possible to prevent distractions and promote safe driving. In this study, we developed a model that could predict the degree of distracted driving based on brain activity. Changes in oxyhemoglobin concentrations were measured in drivers while driving a real car using functional near-infrared spectroscopy (fNIRS). A regression model was constructed for each participant using functional connectivity as an explanatory variable and break reaction time to random beeps while driving as an objective variable. As a result, we were able to construct a prediction model for 11 of the 12 participants. Furthermore, the regression model with the highest prediction accuracy for each participant was analyzed to gain a better understanding of the neural basis of distracted driving. The 11 models were classified into five clusters by hierarchical clustering based on their functional connectivity edges used in each cluster. The results showed that the combinations of the dorsal attention network (DAN)-sensory motor network (SMN) and DAN-ventral attention network (VAN) connections were common in all clusters and that these networks were essential to predict the degree of distraction in complex multitask driving. They also confirmed the existence of multiple types of prediction models with different within- and between-network connectivity patterns. These results indicate that it is possible to predict the degree of distracted driving based on the brain activity of the driver during actual driving. These results are expected to contribute to the development of safe driving systems and elucidation of the neural basis of distracted driving.

## 1. Introduction

Distracted driving is a major cause of traffic accidents; therefore, advanced driving assistance systems and autonomous driving technologies are rapidly being developed. One of the best approaches to preventing drivers from being distracted is to monitor the driver’s attentional state and predict the distraction before it occurs. Some studies have shown that distracted driving is caused by mind wandering during car driving (He et al., 2011; Lohani et al., 2019). The relationship between mental workload, cognitive demands, and the frequency of mind wandering has also been investigated. Mind wandering is a state in which we fall into unconsciously; it is difficult to control and is known to reduce attention and cognitive function. Mind wandering has been studied from a cognitive neuroscience perspective owing to its strong association with cognitive function. Notably, it has also been revealed that decreased attention and cognitive function during driving are reflected in brain activity (Lin et al., 2016; Baldwin et al., 2017; Liu et al., 2017; Xu et al., 2017a, 2017b; Bruno et al., 2018).

Here, we aim to predict driver distraction using preceding brain activity in actual car-driving situations. Few studies have measured brain activity during actual vehicle driving despite the large differences in drivers’ attention and cognitive demands between actual car driving and laboratory experiments using a driving simulator (Jeong et al., 2006; Oka et al., 2015). Yoshino et al. (2013) investigated the relationship between prefrontal cortex activation and actual driving operations on expressways using functional near-infrared spectroscopy (fNIRS) measurements. Their study proved that fNIRS is a powerful technique for robustly measuring brain activity during actual car driving. It remains to be clarified whether it is possible to estimate the degree of distracted driving from the brain activity during actual car driving.

We developed a predictive model of distracted driving based on brain activity measured by fNIRS. A car driving experiment was performed that induced driver distraction and measured brain activity and driving operations, such as braking/accelerator pedaling. Using these measurements, we constructed a regression model using brain activity patterns as the explanatory variable and the driver’s behavioral measure calculated from the driving operations as the objective variable. To characterize brain activity patterns, we used functional connectivity, which is a temporal correlation of brain activity between different measurement locations, as a feature vector. There are several advantages of using functional connectivity, including the following: (1) it can successfully predict individual attentional and psychological states (Shen et al., 2017; Yoo et al., 2018; Cai et al., 2020), and (2) the results of many existing studies in the cognitive neuroscience field can be used to interpret the results of the present study. We investigated the usefulness of using brain activity to estimate the degree of distracted driving. Furthermore, the interpretation of the regression model from the cognitive neuroscience perspective is expected to enrich our understanding of the neural basis of distracted driving. These findings contribute significantly to the development of safe driving systems.

## 2. Material and Method

### 2.1 Participants

Twelve young healthy males (aged 23.4 ± 1.3 years, with driver licenses, and one left-handed individual) participated in this experiment. All participants were informed about the experimental method as well as the risks and signed written informed consent forms. Subsequently, they were required to drive the experimental car along the prescribed course counterclockwise for 10 laps before the experiment. This study was conducted in accordance with the research ethics committee of the Doshisha University, Kyoto, Japan (approval code: 19018). This work was supported by MIC/SCOPE #192297002 and JSPS KAKENHI, Grant Number JP19K12145.

### 2.2 Experimental Design

Car driving is a complex cognitive task that requires attention to many incoming objects while operating multiple components of the driving system such as the steering wheel and pedals. In fact, mind wandering during driving has been confirmed to influence impaired reaction time to emergencies (Yanko and Spalek, 2013). In conventional studies, the driver’s reaction time to randomly occurring events has been used as an attentional index during driving (Yanko and Spalek, 2013; Lin et al., 2016). In this study, distracted driving was induced by forcing drivers to drive on a predetermined simple course for a certain period. In addition, beep tones were presented at random intervals while driving, and the driver was asked to brake in response to each of them. The reaction time to beep tones was measured and used as a behavioral index of distracted driving. Here, we assumed that a shorter the reaction time, indicated a more focused driver, while a slower reaction time, indicated a more distracted driver. Brain activity was measured throughout the experiment and was used to predict reaction time.

The experimental driving course was oval shaped, 40 m long, and flat. In the experiment, the participants were instructed to drive around the course for 15 min while maintaining a vehicle speed of 20 km/h. The beep tones were presented at random intervals within 20-40 seconds from an audio speaker installed behind the driver’s seat, and participants were instructed to brake as soon as possible after recognizing the beeps and to slow down until the vehicle speed reached 10 km/h. The experimental design is illustrated in Figure 1.

**Figure 1.**
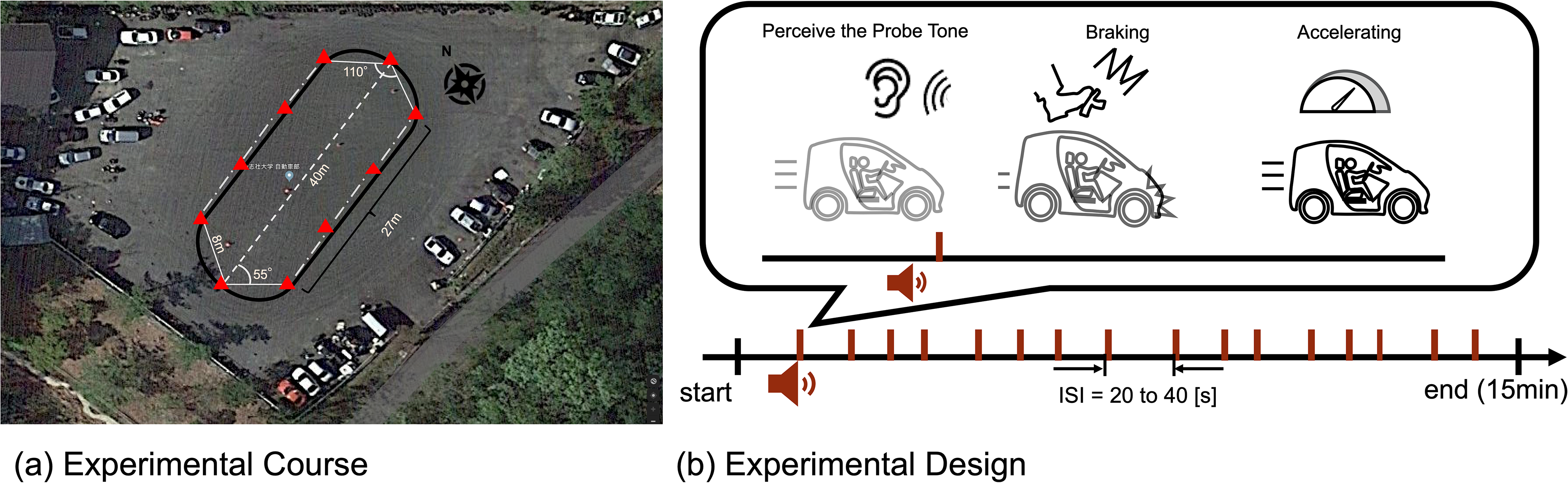
Experimental driving course and design. (A) The experimental course was oval, flat and 40 m long. The red triangles in the figure indicate the safety cones placed along the course. (B) Beeping sounds were generated at random intervals during the 15-minute experiment (inter-stimulus interval: 20-40 s). Participants were required to quickly brake and decelerate to 10 km/h speed if they recognized the beeping sound while traveling at 20 km/h. After decelerating, they accelerated again and were allowed to continue driving while maintaining a speed at 20 km/h.

### 2.3 Data Acquisition

During the experiment, changes in the cerebral blood flow, electrocardiogram, and eye gaze were measured as biometric information, and the brake and accelerator pedal depression and steering angle were measured as indicators of driving operation and of the driver’s state. In this study, a prediction model for distracted driving was developed using changes in cerebral blood flow and brake pedal depression. Other data were collected to be used in future studies and were excluded from the current study. Changes in cerebral blood flow were measured using fNIRS (NIRSport2; NIRx Medical Technologies, Minnesota, USA). The fNIRS is useful for assessing driver performance (Lohani et al., 2019). In this study, the sampling rate was 4.36 Hz and the near-infrared light wavelengths were 730 mm and 810 mm. We used short-distance detector (SDD) probes in addition to conventional source/detector probes to measure and remove physiological noise unrelated to brain activity, such as skin blood flow. Sixteen source probes, 14 detector probes, and 16 SDD probes were arranged as shown in Figure 2. The probe configuration was designed to cover as many brain regions as possible, including the regions involved in attention and motor control associated with driving. A three-dimensional magnetic space digitizer (FASTRAK; Polhemus, Colchester, VT, USA) was used to obtain the coordinates of all probe positions and anatomical landmark positions (Cz, Nz, Iz, and right and left pre-auricular points) for each participant after the driving experiment. The ‘register2polhemus’ function of the NIRS Toolbox (https://github.com/huppertt/nirs-toolbox) was used to determine the Montreal Neurological Institute (MNI) coordinates of the probe positions and estimate those of the measured positions. Moreover, the ‘depthmap’ function of the NIRS Toolbox was used to estimate the brain region corresponding to each measuring channel. Brain regions were determined based on the automated anatomical labeling (AAL) atlas (summarized in Supplementary Table 1). The pupil diameter and eye center coordinates were measured using an ophthalmic eye tracker (Pupil Core monocular, Pupil Labs, Germany), and the ECG was measured using a biological signal amplifier (g.USBamp, g.tec medical engineering GmbH, Austria). A small single-seater electric vehicle (COMS; Toyota Auto Body Co., Ltd.) was used as the experimental vehicle. The vehicle measurement system developed by ITS21 KIKAKU Co., Ltd. (Kanagawa, Japan) was used to record vehicle information, such as brake and accelerator pedal positions, while the steering angle was recorded and shared across multiple computers via a controller area network (CAN). Additionally, a lab-streaming layer (LSL) (https://labstreaminglayer.readthedocs.io/) was used for the unified collection of the multiple measurement time series.

**Figure 2.**
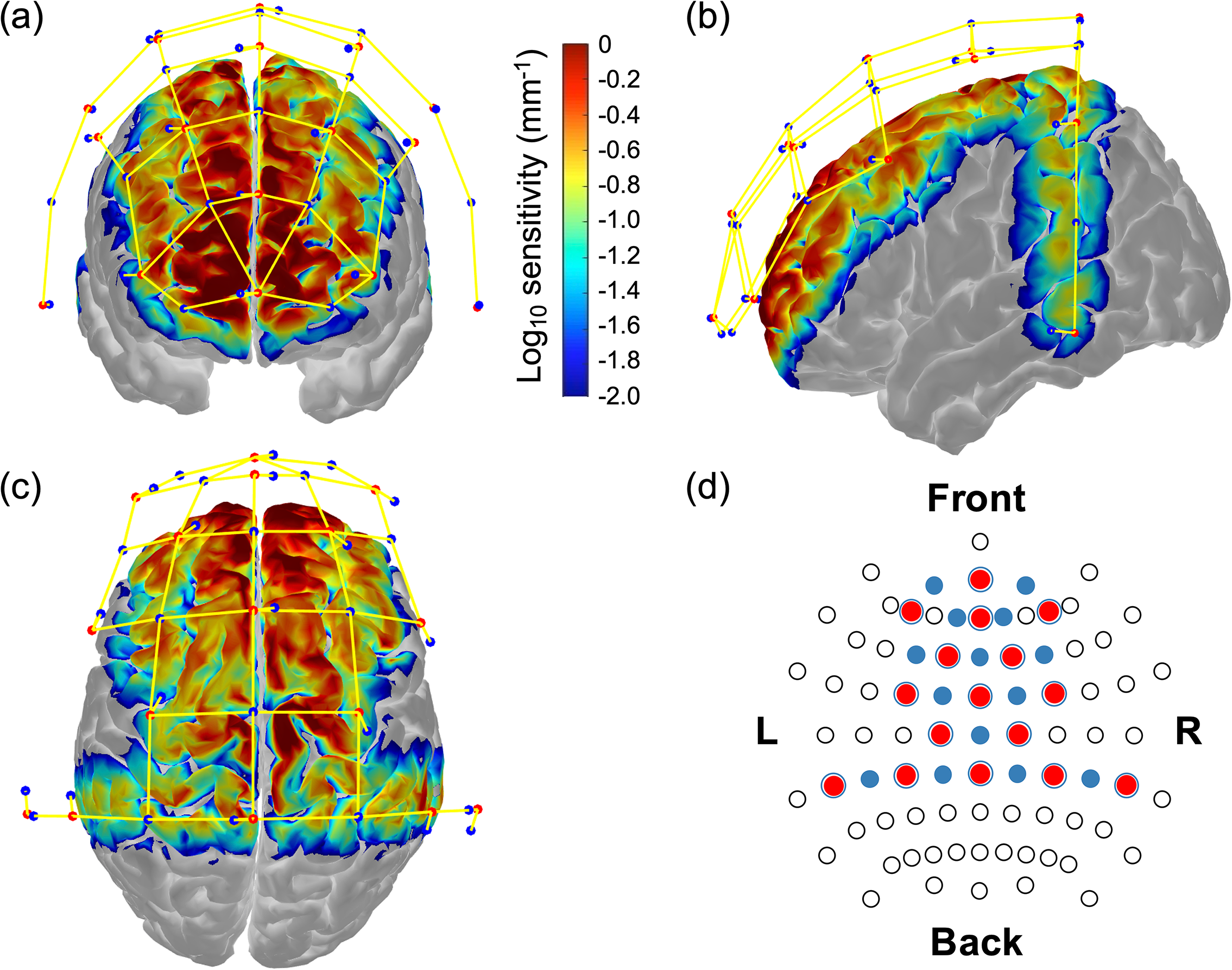
Monte-Carlo simulation results over frontal cortex (A), temporal cortex (B), and parietal cortex (C). Red dots represent emitters, blue dots represent detectors, and yellow lines represent measurement channels, respectively (created using Homer 2 AtlasViewer; v2.8, p2.1: https://www.nitrc.org/frs/shownotes.php?release_id=3956). The color bar represents the spatial sensitivity of fNIRS measurements. The two-dimensional fNIRS montage using International 10-20 measurement system as reference is presented in (D). Blue circles represent short range photosensitive area.

### 2.4 Data analysis

#### 2.4.1 Behavioral data analyses

To assess the degree of distracted driving behavior, we defined the brake reaction times (BRT) from the beep tone to the time when the participant stepped on the brake. To eliminate braking unrelated to the reaction to the beep tones, those that elapsed more than 3 s from the beep tones were ignored. Outlier detection was performed on the calculated BRTs for each participant using the standard deviation method (Ahn et al., 2016). The threshold was set at 3 standard deviations.

#### 2.4.2 fNIRS data analysis: Pre-processing

The fNIRS data were preprocessed using the NIRS Toolbox in MATLAB R2020a (MathWorks Inc., Natick, USA). The following steps were performed to remove physiological noise, such as skin blood flow, heart rate, and pulse, and to convert the measured data into changes in hemoglobin concentration. After converting the raw intensity signals to changes in optical density, the optical density values were converted into oxy- and deoxygenated hemoglobin concentration changes using the modified Beer-Lambert law. To remove physiological noise in the whole body, bandpass filtering was performed with passband of 0.01 Hz and 0.1 Hz (Yoshino et al., 2013; Ahn et al., 2016). Finally, short separation regression (SSR) was applied to suppress the effects of skin blood flow (Yücel et al., 2015; Tachtsidis and Scholkmann, 2016).

#### 2.4.3 fNIRS Data Analysis: Functional connectivity

Functional connectivity (FC) is a measure of the temporal synchronization between two brain activities. The Pearson’s correlation coefficient was used to calculate the FC. The FC was calculated for the oxy-Hb time series before the beep-tone presentation. The oxy-Hb data were extracted using different window sizes of 10, 15, and 20 s. We chose 10 s because the lower cutoff frequency of the bandpass filtering was 0.01 Hz and 20 s, as the maximum inter-stimulus interval was set to 20 s. Because our fNIRS measurement was performed on 44 channels, a 44 × 44 FC matrix was computed for each beep-tone presentation. The calculated FC matrix was then Fisher Z-transformed (Douw et al., 2016) to improve the normality of the correlation coefficients (Lu et al., 2019). Because the FC matrix was symmetric, 964-dimensional FC elements (triangular parts of the FC matrix) were vectorized and used as explanatory variables for the regression.

### 2.5 Proposed methods

An overview of our proposed method for building a prediction model for distracted driving is shown in Figure 3. We constructed a regression model using the functional connectivity (946-dimensional vector) of the preceding brain activity before the beep tone was presented as the explanatory variable, and the corresponding BRT as the objective variable. The model was constructed for each participant using a dataset of multiple trials (braking reaction). The detailed process of each step is described below.

**Figure 3.**
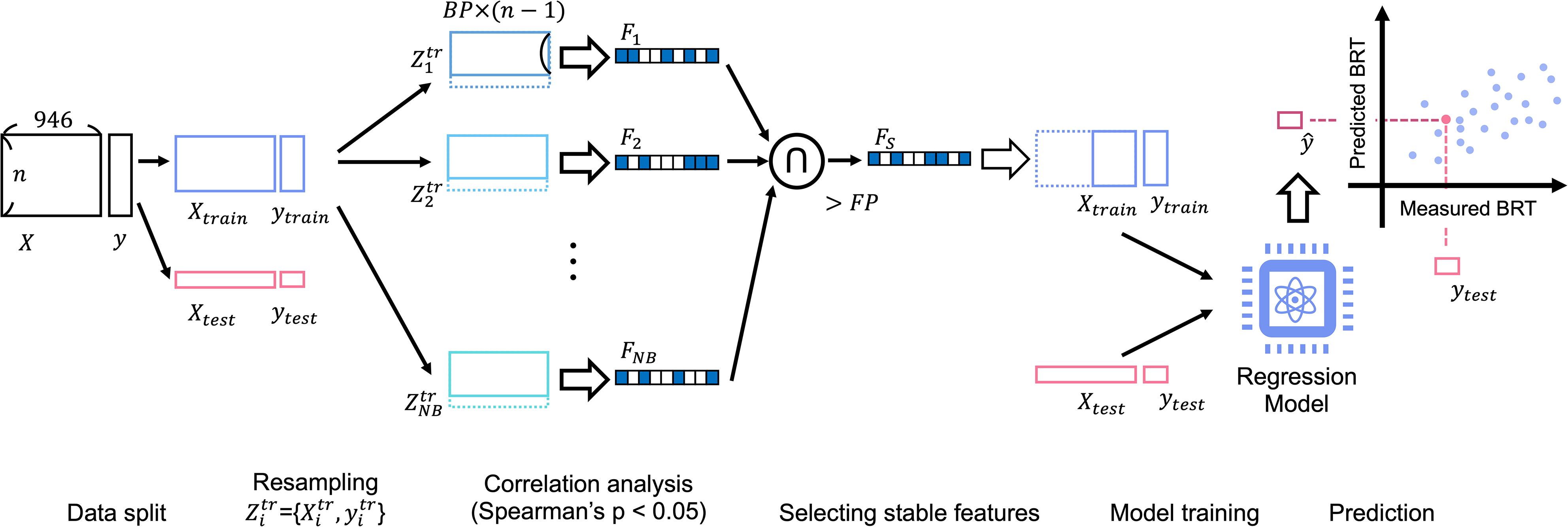
The overview of our proposed method to build prediction model of the distracted driving. Cross-validation and feature selection were incorporated into the model building process. The model was constructed for each participant using the dataset from multiple car braking trials. NB: number of resampling, BP: proportion of sampling, FP: stability of the features.

### 2.6 Feature selection: Resampling and correlation analysis

Feature selection was performed to reduce the dimensionality of FC data. In this study, we used a bootstrap-based feature selection method (Wei et al., 2020). The goal of this method is to identify stable features that can be consistently identified among resampled subsets. We defined the hyperparameters of the method: number of bootstrapping (NB) as the sampling times, bootstrap percentage (BP) as the proportion of sampling, and frequency percentage (FP) as the stability of the features used to determine the final feature sets in the NB subsets. For example, BP = 50% indicated that half of all trials (braking operations) were chosen from the original dataset to form a subset. In addition, FP = 50% indicated that the corresponding feature was chosen for 50% of the NB subsets.

We sampled the training data for NB times without replacement for the number of samples multiplied by BP, as shown in Figure 3. Spearman’s correlation coefficients comparing each feature (FC element) and the behavioral data (BRT) in the sampled subset were calculated, and significant features, having p-values lower than 0.05, were chosen as the promising features. After completing the correlation analysis for all subsets, the features where the FP was higher than the threshold among the NB subsets were chosen as the final features and used for model construction.

### 2.7 Model training and evaluation

To link brain activity to the degree of side-driving, a linear regression model was constructed with FC as the explanatory variable and BRT as the objective variable. Ordinary least squares (OLS) regression was used to estimate the coefficients of the linear regression model; the OLS algorithm minimized the squared sum of the residuals. Here, we used *fitrlinear* function implemented in MATLAB 2020a. In the regression model, the lower triangular part of each FC matrix was vectorized and the corresponding BRT was used as the objective variable. The data matrices for the explanatory and objective variables consisted of *N* × 964 and *N* × 1, respectively, where *N* was the number of breaking responses for each participant.

The model training process included leave-one-out cross-validation and feature selection, as shown in Figure 3. Feature selection and model construction were repeatedly performed using N-1 samples and the built model was validated using each excluded sample. Finally, the N pairs of the predicted and measured BRT data were obtained and used to calculate Spearman’s rank correlation as a measure of the overall prediction performance (test of no correlation; p < 0.05). The aforementioned model training and evaluation were performed for each participant. Furthermore, we used different hyperparameter settings for bootstrapping (NB = 10, 20, 50, 100, 300, 500, and 1000; BP = 60 and 80%; FP = 50, 60, 70, 80, 90, and 100%) and window sizes (10, 15, and 20 s), and determined the optimal setting that maximized the prediction accuracy for each participant.

### 2.8 Interpreting the individual regression model

To gain a better understanding of the neural basis of distracted driving, we analyzed a regression model that determined the best prediction accuracy for each participant. Previous studies have shown that it is essential to consider individual differences when predicting mind wandering while driving (Zhang and Kumada, 2018). To capture features unique to each participant and those that were similar across individuals, we performed data clustering to classify the patterns of FC edges selected by feature selection in the model construction of the individual models.

In the clustering analysis, the selected patterns of FC edges were represented by a matrix showing the connectivity between 116 regions defined by the AAL atlas, where the elements of the matrix indicated whether the connectivity between two regions was selected by a feature selection, with a value of 1 or 0. The Jaccard distance was used as the distance measure between the two samples, and the patterns of FC edges used in the individual regression models were classified by agglomerative hierarchical clustering using the linkage function implemented in MATLAB R2020a. Here, the unweighted pair group method with arithmetic mean (UPGMA) was chosen for distance calculation. Finally, the similarity of the model features for each individual was visualized in a phylogenetic tree by applying the dendrogram function in MATLAB R2020a. The common features within each cluster were interpreted in terms of the well-known functional network to which they belonged and how they differed between clusters.

## 3. Result

### 3.1 Regression model for each individual

The accuracy of each of the 12 individual regression models is summarized in Figure 4, together with the connectome plots consisting of the FC edges chosen via feature selection (drawn using BrainNet Viewer: https://www.nitrc.org/projects/bnv/). We successfully constructed a model with sufficient prediction accuracy (p < 0.05) for 11 of 12 participants. The figure includes a bar chart indicating the correlation between the measured and predicted BRTs for each individual model. The results indicated that the regression model differed between individuals in their chosen FC edges and number of edges.

**Figure 4.**
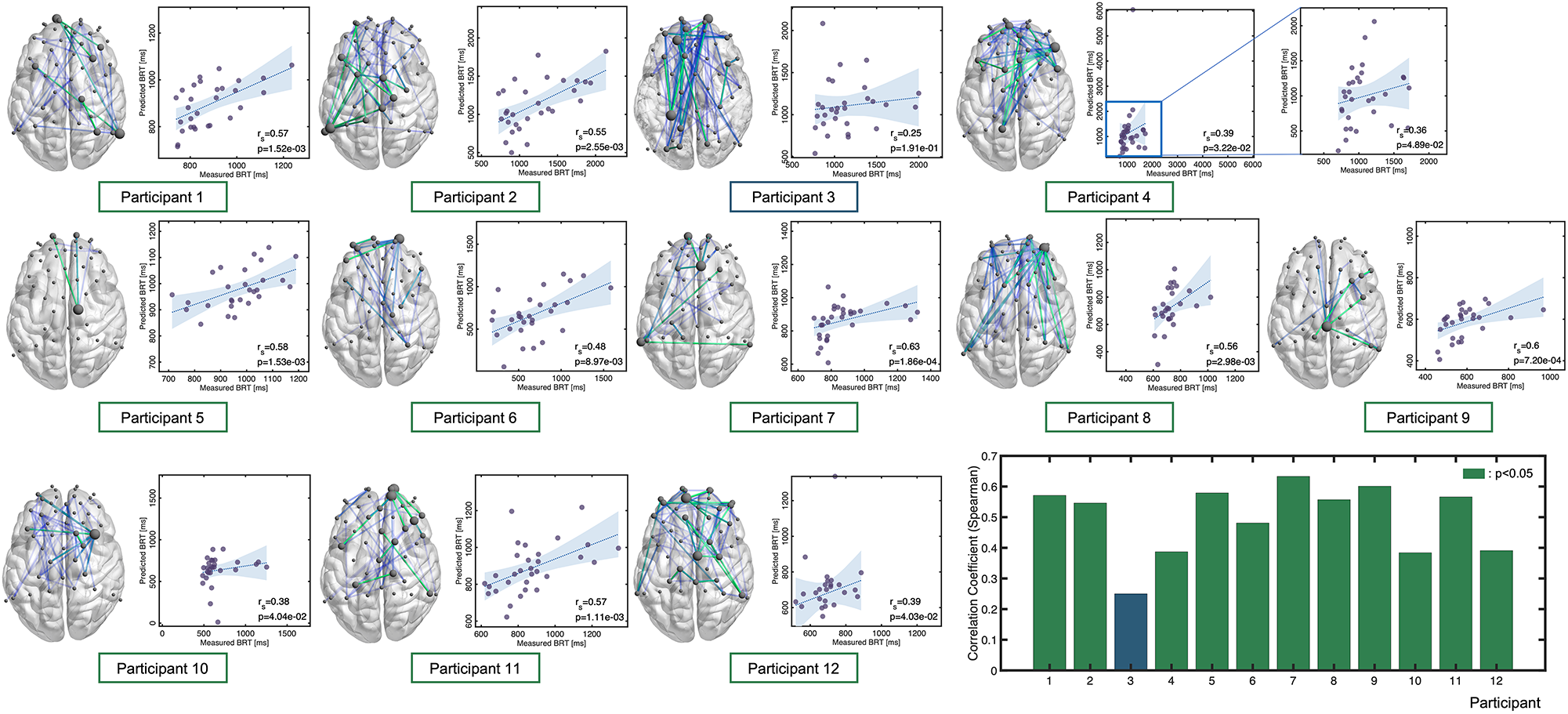
Correlation between the predicted and measured BRTs for each participant. Each connectome plot indicates the functional connectivity edges chosen via feature selection and used for the regression model. The bar graph summarizes the Spearman’s correlation as accuracy measure for each individual model.

The distribution of the measured the BRT for each participant is summarized in Figure 5, which indicated that the distribution of the BRT differed among the participants. Outliers in the BRT were identified in subjects 4, 5, 8, 9, and 12, which are indicated by red dots in Figure 5. They were removed for the regression analysis for each participant.

**Figure 5.**
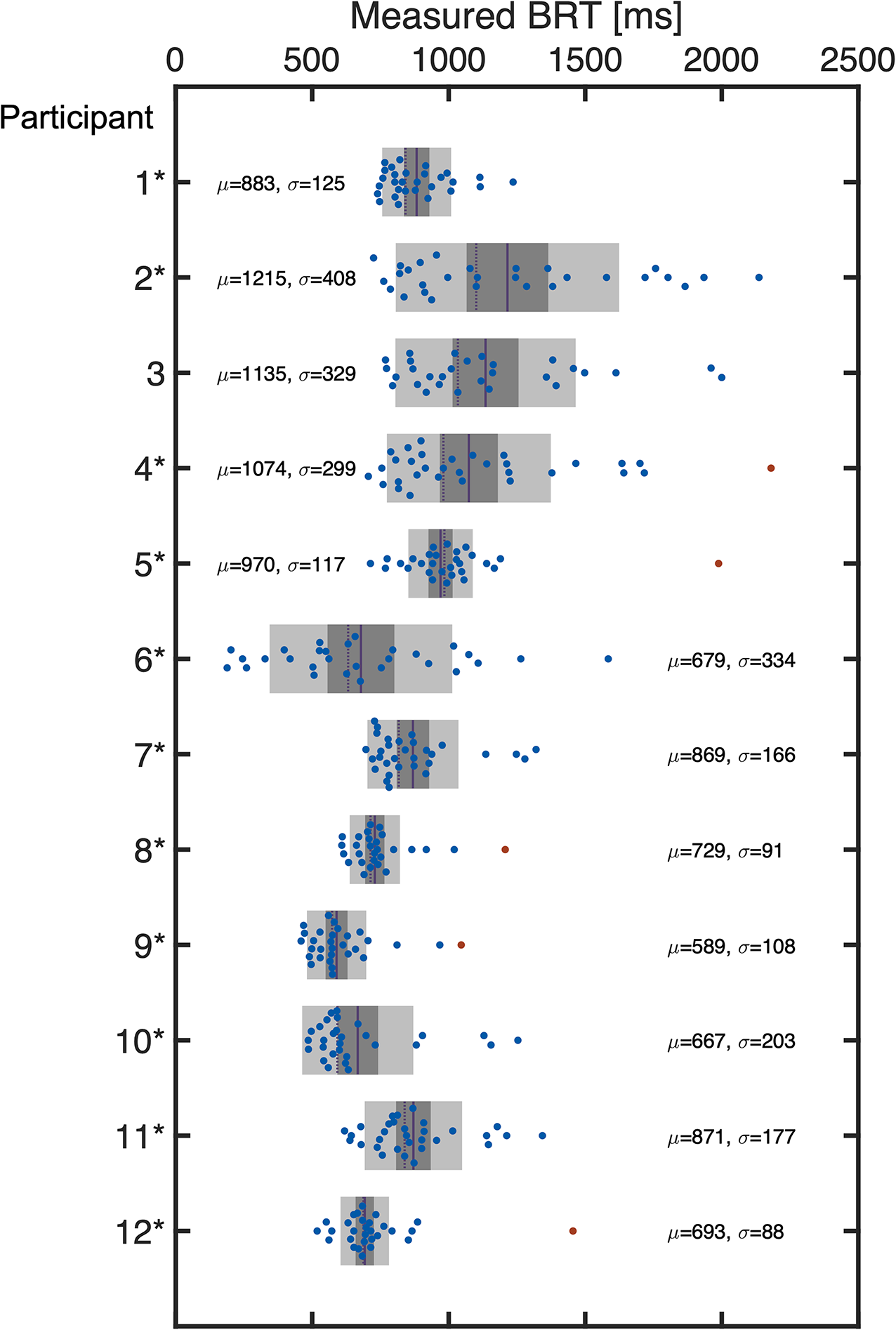
Distribution of BRT measured for each participant. The red dots indicate the outliers excluded for the regression modeling. The mean (μ) and standard deviation (σ) are also indicated for each participant.

Accuracy and the optimized hyperparameters are summarized in Table 1, which also summarizes the number of samples (i.e., the number of breaking reactions, *N*) for each participant and the number of FC edges selected in feature selection averaged over the cross-validation folds. The results showed that the feature selection successfully reduced the dimension of the explanatory variables from 946 to a number below that of the sample size.

**Table 1.**
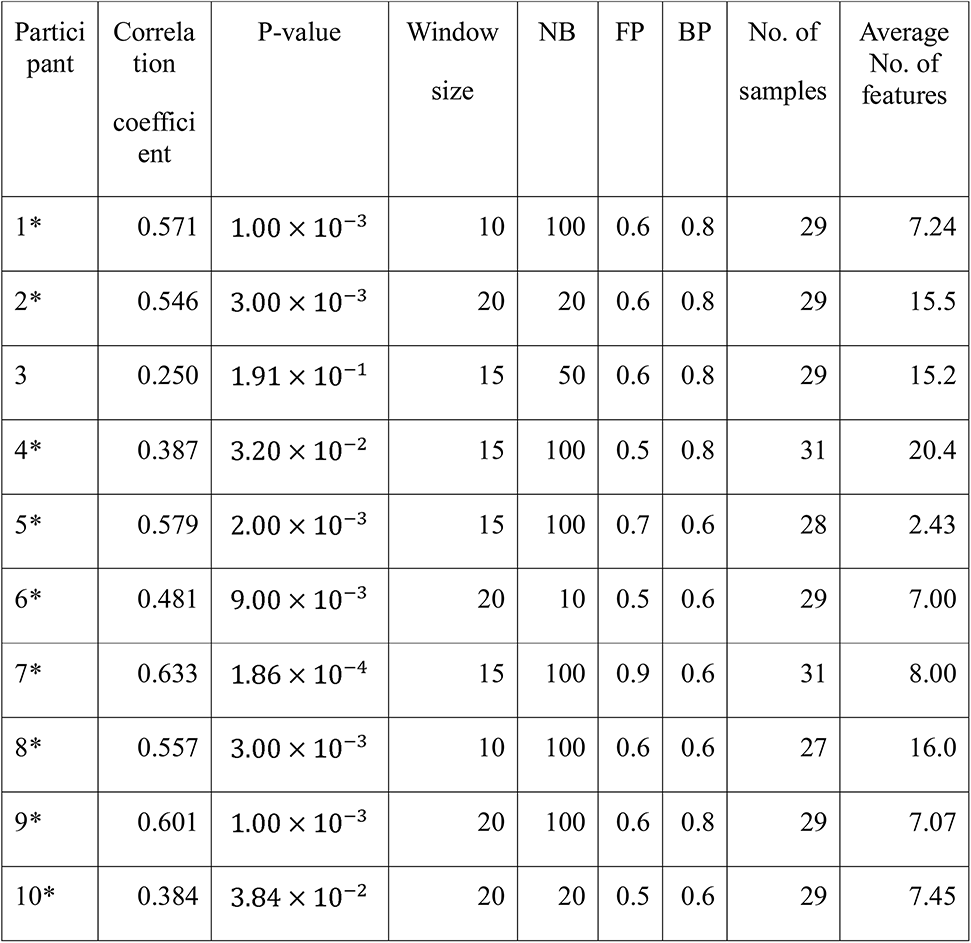
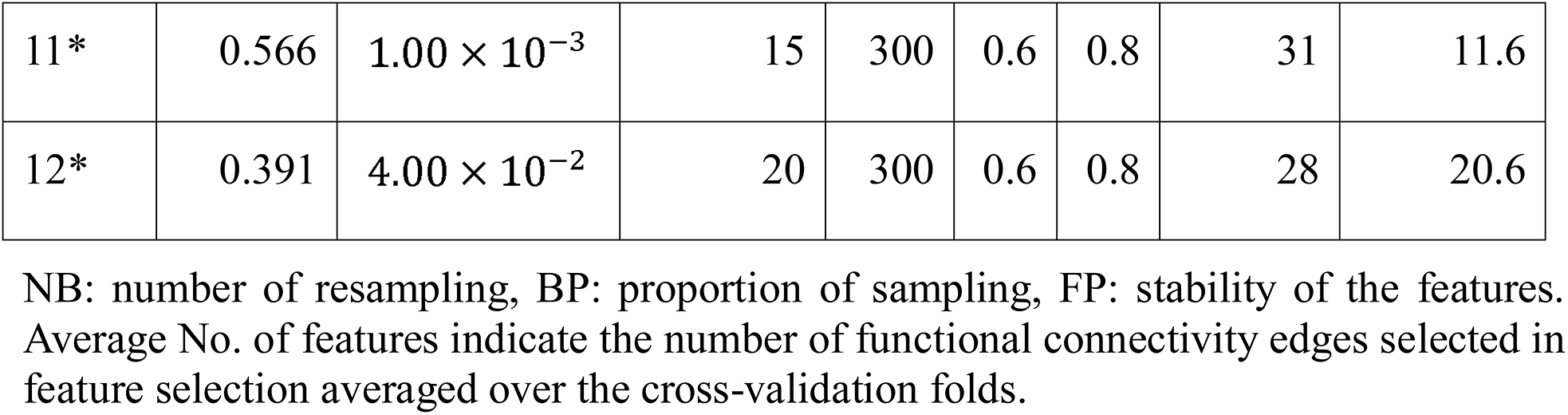
The prediction results and the optimized hyperparameters for the individual regression models (*: p < 0.05).

### 3.2 Interpreting individual regression models

Eleven of the 12 individuals, whose regression models indicated sufficient accuracy, were classified by hierarchical clustering based on the similarity of the selected FC edges of each individual model. Figure 6 shows a dendrogram of the FC patterns of the 11 individual models. Through clustering, the 11 individual models were classified into 5 clusters (the red line drawn on the dendrogram indicates the distance threshold used for splitting into 5 clusters). Commonly selected FC edges in clusters are shown in the connectome plot. The connectome is also represented as a circle plot. The color of the node in the circle plot indicates which of the six well-known brain functional networks (dorsal attention network: DAN, default mode network: DMN, sensorimotor network: SMN, ventral attention network: VAN, frontoparietal network: FPN, and salience network: SN) the brain region comprising that FC edge belongs to. The results indicated that although the FC patterns differed between clusters, there were some common inter- and intra-network connections between clusters.

**Figure 6.**
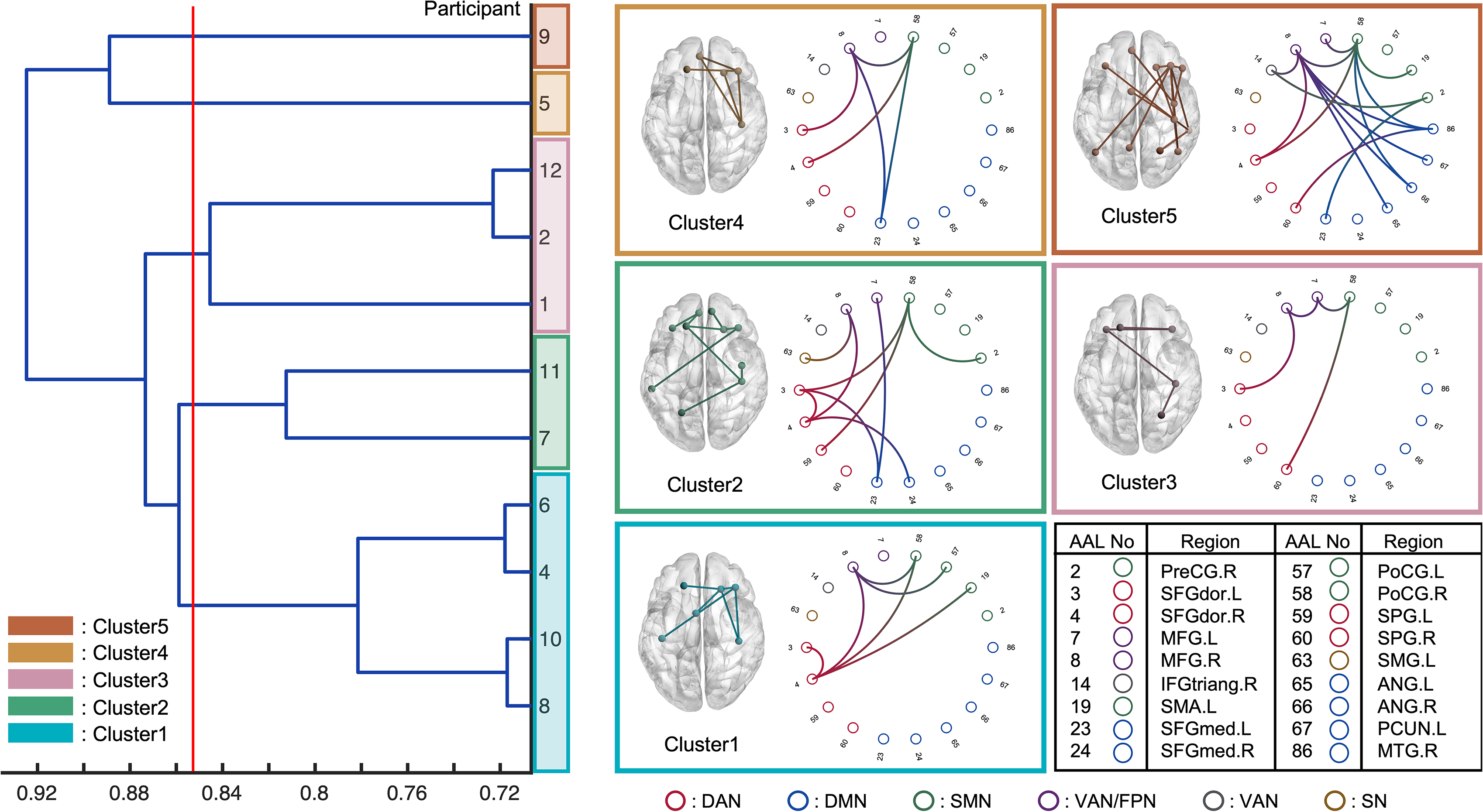
Eleven individual models with the sufficient prediction accuracy were classified using a hierarchical clustering based on the functional connectivity chosen via feature selection. The horizontal axis indicates the Jaccard distance among the models. The red line was drawn to analyze the five clusters. The circle plot and connectome indicate the common functional connectivity edges within each cluster. Each node channel was associated with brain region determined by automated anatomical labeling (AAL) atlas. The nodes are colored based on which well-known functional network they belonged to: DAN, dorsal attention network; DMN, default mode network; SMN, Sensorimotor network; VAN, ventral attention network; FPN, frontoparietal network; SN, salience network.

## 4. Discussion

### 4.1 Performance of the prediction model

We constructed a regression model with significant prediction accuracy for 11 of the 12 participants. Of note, participant 3 did not achieve sufficient accuracy. The BRT distribution of each participant is indicated in Figure 5, and for Participant 3, outliers were not detected even though there was less data around 2000 ms. An imbalance in the dataset is problematic in regression modeling because it leads to an imbalance in the training data for each CV fold. Improvement of the outlier detection scheme could address this issue. Nonetheless, our results suggest that fNIRS functional connectivity can be used to predict the variability of braking response during car driving and can be utilized as an indicator of distracted driving.

The hyperparameter settings for each model were determined. The parameters for feature selection, NB, FP, and BP, differed among individuals, as shown in Table 1. A variability that has also been confirmed in previous studies that proposed bootstrap-based feature selection (Wei et al., 2020), while the effectiveness of this scheme was confirmed for our fNIRS functional connectivity-based prediction model. Furthermore, different window sizes were adopted for the functional connectivity analysis for each participant. It was reasonable to choose different window sizes for each individual because functional connectivity varies across individuals and multiple timescales, as revealed by Telesford et al. (2016).

### 4.2 Interpretation of the prediction model

The interpretation of the constructed model is expected to elucidate the neural basis of distracted driving. For example, the FC edges identified, which are commonly employed across individuals, suggest that these play an important role in predicting the degree of distracted driving. In addition, if the choices of FC edges in the individual models can be classified into several patterns, the common FC edges within each cluster will provide representative patterns of the FC network associated with distracted driving. Herein, we interpreted the predictive models in terms of which FC edges were used as the explanatory variables and which well-known functional networks (DAN, DMN, SMN, VAN, FPN, and SN). The characteristics of each cluster are summarized as the functional networks contributed to in Figure 6.

First, we found a common between-network connectivity of DAN-SMN and DAN-VAN across all clusters. DAN plays an important role in top-down attentional control (Taren et al., 2017), which is essential for driving. In addition, a previous study that predicted multitasking performance using functional connectivity at rest reported that DAN-SMN connectivity during a task can predict multitasking performance (Wen et al., 2018). Car driving requires complex multitasking, which is essential in driving experiments. These previous findings support the idea that the DAN-SMN connection is necessary for predicting the degree of distracted driving.

For between-network connectivity of the DAN-VAN, the connectivity between the middle frontal gyrus (MFG) and dorsolateral superior frontal gyrus (SFGdor) was confirmed in all clusters. The MFG is a node region of the VAN that is involved in bottom-up attention (Doricchi et al., 2010) and attentional control (Japee et al., 2015). In addition, the left dorsolateral prefrontal cortex (DLPFC) including SFGdor and MFG have been reported to play an important role in spatial recognition in a previous study used the working memory task (Ren et al., 2019). Because car driving involves paying attention to multiple objects in a driver’s field of view, it is convincing that the FC edges, SFGdor and MFG, were used for regression modeling. In a previous study on meditation, a technique indicated as an effective approach for reducing mind wandering, the increase in FC between the SFGdor.R and MFG.R was observed when the meditators attempted to focus their attention on their main task from task-unrelated thought (He et al., 2021). This result was also confirmed by an fMRI experiment (Hasenkamp et al., 2012). These previous findings also supported our results because the drivers were expected to shift their attention back from distractors to the driving task when their minds were distracted. In addition, Lin et al. (2016) in their electroencephalography-based driving experiments, driving was essentially a complex continuous tracking task requiring bottom-up processes for the multisensory integration of information from the external environment, as well as top-down modulatory influences based on the driver’s internal goals, strategies, and current intentions. Therefore, it is reasonable that DAN-VAN connections are essential for predicting the degree of distracted driving.

Next, we discuss the characteristic within each of these five clusters. Cluster-specific between-network connectivity was not observed in Clusters 1 and 3. These two clusters mainly consisted of DAN-VAN and DAN-SMN, which were commonly observed across all clusters. Cluster-specific FC edges were found in SFGdor.L-SFGdor.R for cluster 1 and MFG.L-MFG.R for Cluster 3. Since each of these FC edges was within-network connectivity of the DAN and VAN, Clusters 1 and 3 were the same in the patterns of functional networks chosen for predictive modeling, except for their use of within-network connectivity. It can be concluded that Cluster 1 was DAN-weighted, while Cluster 3 was a VAN weighted model (individual). DAN and VAN are involved in top-down (Taren et al., 2017) and bottom-up attention, (Doricchi et al., 2010) respectively, and both were essential for car driving tasks.

In Cluster 2, cluster-specific between-network connectivity of the DAN-DMN was also SFGdor.L-SFGmed.L and SFGdor.R-SFGmed.R connectivity. The medial prefrontal cortex (mPFC), including SFGmed.R, is one of the key regions of the DMN (Vatansever et al., 2015). Yamashita et al. (2021) revealed that there were two dominant brain states during sustained attention: one characterized by DMN and limbic network activation, and the other by activation of DAN, SMN, salience, and visual networks. Referring to their findings, it is convincing that the DAN-DMN connectivity was chosen for the model in Cluster 2. In addition, the between-network connectivity of the FPN-DMN and DAN-FPN was found to be MFG.L-SFGmed.R and SFGdor.R-MFG.R connectivity. The MFG is included in the FPN (Zhu et al., 2017) as well as the VAN. Dixon et al. (2018) identified two subsystems of the FPN: one had stronger connectivity with the DMN than with the DAN, and the other had the opposite connectivity. In addition, Spreng et al. (2013) stated that the FPN acts as a "gatekeeper" that regulates the DMN, which is responsible for internal cognition, and the DAN, which is responsible for external cognition. Our results in Cluster 2 were consistent with their ideas, and we concluded that Cluster 2 was the DMN-and DAN-based model gated with FPN.

In Cluster 4, SFGmed.L-PoCG.R connectivity was found to be a specific FC and differed from that of other clusters. The PoCG is included in the SMN (Cui et al., 2020), and the mPFC, including the SFGmed, has been reported to not only to project to the limbic regions but also to connect to the motor system, which is why it plays an important role in motor planning and formation (Jiang et al., 2019). This is consistent with the findings for SFGmed.L-PoCG.R the FC edge was chosen for the prediction of driving behavior in Cluster 4. Thus, we conclude that Cluster 4 was an SMN-based model.

In Cluster 5, as in Cluster 2, the between-network connectivity of the FPN-DMN and FPN-DAN was used for modeling. Furthermore, the SFGmed.L-PreCG.R connectivity was found to be a cluster-specific connectivity. However, it could be regarded as the same as Cluster 4 in terms of PreCG being included in the SMN. These observations imply that Cluster 5 was a mixed model of Clusters 2 and 4.

In this study, we used regression modeling to predict the degree of distracted driving from FC for each participant, using brain activity measured in an actual car driving environment. We successfully constructed predictive models for 11 of the 12 participants. In addition, to analyze the features common to all participants and features specific to individual patterns, the 11 models that showed sufficient prediction accuracy were classified into five patterns based on the patterns of FC edges extracted in feature selection during model construction. The results showed that the combination of DAN-SMN and DAN-VAN networks was common to all clusters, indicating that these networks were essential for predicting the degree of distraction in driving a car, which is a complex multitask behavior. The individual features of each of the five clusters were also interpreted as follows: a model with within-network connectivity of DAN and VAN was added to the underlying DAN-SMN and DAN-VAN FCs, respectively (Clusters 1 and 3); and a model based on DMN and DAN, gated by FPN (Cluster 2) and an SMN-based model (Cluster 4); and a mixture of Clusters 2 and 4 (Cluster 5) were constructed. Referring to the results of previous studies, all five models were relevant for driving behavior and distracted driving. Our results will contribute to the development of brain activity-based systems for predicting distracted driving to promote safe driving as well as to the understanding of the neural basis of distracted driving.

## 5. Abbreviations

AAL: Automated anatomical labeling
BP: Bootstrap percentage
BRT: Brake reaction times
CAN: Controller area network
DAN: Dorsal attention network
FC: Functional connectivity
FP: Frequency percentage
LSL: Lab-streaming layer
MFG: Middle frontal gyrus
MNI: Montreal Neurological Institute
NB: Number of bootstrapping
OLS: Ordinary least squares
SDD: Short-distance detector
SMN: Sensory motor network
SN: Salience network
SSR: Short separation regression
UPGMA: Unweighted pair group method with arithmetic
VAN: Ventral attention network
DLPFC: Dorsolateral prefrontal cortex
DMN: Default mode network
ECG: Electrocardiography
FPN: frontoparietal network
NIRS: Near-infrared spectroscopy

## 6. Data Availability Statement

The datasets generated for this study are available on request to the corresponding author.

## 7. Ethics Statement

Studies involving human participants were reviewed and approved by the Research Ethics Committee of Doshisha University, Kyoto, Japan (approval code: 19018). The individuals provided written informed consent to participate in the study.

## 10. Acknowledgement

We would like to thank Editage (www.editage.com) for English language editing.

## References

Ahn, S., Nguyen, T., Jang, H., Kim, J. G., and Jun, S. C. (2016). Exploring Neuro-Physiological Correlates of Drivers’ Mental Fatigue Caused by Sleep Deprivation Using Simultaneous EEG, ECG, and fNIRS Data. Frontiers in Human Neuroscience 0, 219. doi:10.3389/FNHUM.2016.00219.

Baldwin, C. L., Roberts, D. M., Barragan, D., Lee, J. D., Lerner, N., and Higgins, J. S. (2017). Detecting and Quantifying Mind Wandering during Simulated Driving. Frontiers in Human Neuroscience 0, 406. doi:10.3389/FNHUM.2017.00406.

Bruno, J. L., Baker, J. M., Gundran, A., Harbott, L. K., Stuart, Z., Piccirilli, A. M., et al. (2018). Mind over motor mapping: Driver response to changing vehicle dynamics. Human Brain Mapping 39, 3915–3927. doi:10.1002/HBM.24220.

Cai, H., Chen, J., Liu, S., Zhu, J., and Yu, Y. (2020). Brain functional connectome-based prediction of individual decision impulsivity. Cortex 125, 288–298. doi:10.1016/J.CORTEX.2020.01.022.

Cui, G., Ou, Y., Chen, Y., Lv, D., Jia, C., Zhong, Z., et al. (2020). Altered Global Brain Functional Connectivity in Drug-Naive Patients With Obsessive-Compulsive Disorder. Frontiers in Psychiatry 11, 98. doi:10.3389/FPSYT.2020.00098/BIBTEX.

Dixon, M. L., de La Vega, A., Mills, C., Andrews-Hanna, J., Spreng, R. N., Cole, M. W., et al. (2018). Heterogeneity within the frontoparietal control network and its relationship to the default and dorsal attention networks. Proceedings of the National Academy of Sciences of the United States of America 115, E1598–E1607. doi:10.1073/PNAS.1715766115/-/DCSUPPLEMENTAL.

Doricchi, F., MacCi, E., Silvetti, M., and MacAluso, E. (2010). Neural Correlates of the Spatial and Expectancy Components of Endogenous and Stimulus-Driven Orienting of Attention in the Posner Task. Cerebral Cortex 20, 1574–1585. doi:10.1093/CERCOR/BHP215.

Douw, L., Wakeman, D. G., Tanaka, N., Liu, H., and Stufflebeam, S. M. (2016). State-dependent variability of dynamic functional connectivity between frontoparietal and default networks relates to cognitive flexibility. Neuroscience 339, 12–21. doi:10.1016/J.NEUROSCIENCE.2016.09.034.

Hasenkamp, W., Wilson-Mendenhall, C. D., Duncan, E., and Barsalou, L. W. (2012). Mind wandering and attention during focused meditation: A fine-grained temporal analysis of fluctuating cognitive states. NeuroImage 59, 750–760. doi:10.1016/J.NEUROIMAGE.2011.07.008.

He, H., Hu, L., Zhang, X., and Qiu, J. (2021). Pleasantness of mind wandering is positively associated with focus back effort in daily life: Evidence from resting state fMRI. Brain and Cognition 150, 105731. doi:10.1016/J.BANDC.2021.105731.

He, J., Becic, E., Lee, Y.-C., and McCarley, J. S. (2011). Mind Wandering Behind the Wheel: Performance and Oculomotor Correlates. *https://doi.org/10.1177/0018720810391530* 53, 13–21. doi:10.1177/0018720810391530.

Japee, S., Holiday, K., Satyshur, M. D., Mukai, I., and Ungerleider, L. G. (2015). A role of right middle frontal gyrus in reorienting of attention: A case study. Frontiers in Systems Neuroscience 9, 23. doi:10.3389/FNSYS.2015.00023/ABSTRACT.

Jeong, M., Tashiro, M., Singh, L. N., Yamaguchi, K., Horikawa, E., Miyake, M., et al. (2006). Functional brain mapping of actual car-driving using [18F]FDG-PET. Annals of Nuclear Medicine 2006 20:9 20, 623–628. doi:10.1007/BF02984660.

Jiang, W., Lei, Y., Wei, J., Yang, L., Wei, S., Yin, Q., et al. (2019). Alterations of interhemispheric functional connectivity and degree centrality in cervical dystonia: A resting-state fMRI study. Neural Plasticity 2019. doi:10.1155/2019/7349894.

Lin, C.-T., Chuang, C.-H., Kerick, S., Mullen, T., Jung, T.-P., Ko, L.-W., et al. (2016). Mind-Wandering Tends to Occur under Low Perceptual Demands during Driving. Scientific Reports 6. doi:10.1038/SREP21353.

Liu, Z., Zhang, M., Xu, G., Huo, C., Tan, Q., Li, Z., et al. (2017). Effective Connectivity Analysis of the Brain Network in Drivers during Actual Driving Using Near-Infrared Spectroscopy. Frontiers in Behavioral Neuroscience 0, 211. doi:10.3389/FNBEH.2017.00211.

Lohani, M., Payne, B. R., and Strayer, D. L. (2019). A Review of Psychophysiological Measures to Assess Cognitive States in Real-World Driving. Frontiers in Human Neuroscience 0, 57. doi:10.3389/FNHUM.2019.00057.

Lu, X., Li, T., Xia, Z., Zhu, R., Wang, L., Luo, Y. J., et al. (2019). Connectome-based model predicts individual differences in propensity to trust. Human Brain Mapping 40, 1942–1954. doi:10.1002/HBM.24503.

Oka, N., Yoshino, K., Yamamoto, K., Takahashi, H., Li, S., Sugimachi, T., et al. (2015). Greater Activity in the Frontal Cortex on Left Curves: A Vector-Based fNIRS Study of Left and Right Curve Driving. PLOS ONE 10, e0127594. doi:10.1371/JOURNAL.PONE.0127594.

Ren, Z., Zhang, Y., He, H., Feng, Q., Bi, T., and Qiu, J. (2019). The different brain mechanisms of object and spatial working memory: Voxel-based morphometry and resting-state functional connectivity. Frontiers in Human Neuroscience 13, 248. doi:10.3389/FNHUM.2019.00248/BIBTEX.

Shen, X., Finn, E. S., Scheinost, D., Rosenberg, M. D., Chun, M. M., Papademetris, X., et al. (2017). Using connectome-based predictive modeling to predict individual behavior from brain connectivity. Nature Protocols 2017 12:3 12, 506–518. doi:10.1038/nprot.2016.178.

Spreng, R. N., Sepulcre, J., Turner, G. R., Stevens, W. D., and Schacter, D. L. (2013). Intrinsic architecture underlying the relations among the default, dorsal attention, and frontoparietal control networks of the human brain. Journal of cognitive neuroscience 25, 74–86. doi:10.1162/JOCN_A_00281.

Tachtsidis, I., and Scholkmann, F. (2016). False positives and false negatives in functional near-infrared spectroscopy: issues, challenges, and the way forward. *https://doi.org/10.1117/1.NPh.3.3.031405* 3, 031405. doi:10.1117/1.NPH.3.3.031405.

Taren, A. A., Gianaros, P. J., Greco, C. M., Lindsay, E. K., Fairgrieve, A., Brown, K. W., et al. (2017). Mindfulness Meditation Training and Executive Control Network Resting State Functional Connectivity: A Randomized Controlled Trial. Psychosomatic medicine 79, 674. doi:10.1097/PSY.0000000000000466.

Telesford, Q. K., Lynall, M. E., Vettel, J., Miller, M. B., Grafton, S. T., and Bassett, D. S. (2016). Detection of functional brain network reconfiguration during task-driven cognitive states. NeuroImage 142, 198–210. doi:10.1016/J.NEUROIMAGE.2016.05.078.

Vatansever, D., Menon, D. K., Manktelow, A. E., Sahakian, B. J., and Stamatakis, E. A. (2015). Default mode network connectivity during task execution. NeuroImage 122, 96–104. doi:10.1016/J.NEUROIMAGE.2015.07.053.

Wei, L., Jing, B., and Li, H. (2020). Bootstrapping promotes the RSFC-behavior associations: An application of individual cognitive traits prediction. Human Brain Mapping 41, 2302–2316. doi:10.1002/HBM.24947.

Wen, T., Liu, D. C., and Hsieh, S. (2018). Connectivity patterns in cognitive control networks predict naturalistic multitasking ability. Neuropsychologia 114, 195–202. doi:10.1016/J.NEUROPSYCHOLOGIA.2018.05.002.

Xu, G., Zhang, M., Wang, Y., Liu, Z., Huo, C., Li, Z., et al. (2017a). Functional connectivity analysis of distracted drivers based on the wavelet phase coherence of functional near-infrared spectroscopy signals. PLOS ONE 12, e0188329. doi:10.1371/JOURNAL.PONE.0188329.

Xu, L., Wang, B., Xu, G., Wang, W., Liu, Z., and Li, Z. (2017b). Functional connectivity analysis using fNIRS in healthy subjects during prolonged simulated driving. Neuroscience Letters 640, 21–28. doi:10.1016/J.NEULET.2017.01.018.

Yamashita, A., Rothlein, D., Kucyi, A., Valera, E. M., and Esterman, M. (2021). Brain state-based detection of attentional fluctuations and their modulation. NeuroImage 236, 118072. doi:10.1016/J.NEUROIMAGE.2021.118072.

Yanko, M. R., and Spalek, T. M. (2013). Driving With the Wandering Mind: The Effect That Mind-Wandering Has on Driving Performance. *https://doi.org/10.1177/0018720813495280* 56, 260–269. doi:10.1177/0018720813495280.

Yoo, K., Rosenberg, M. D., Hsu, W. T., Zhang, S., Li, C. S. R., Scheinost, D., et al. (2018). Connectome-based predictive modeling of attention: Comparing different functional connectivity features and prediction methods across datasets. NeuroImage 167, 11–22. doi:10.1016/J.NEUROIMAGE.2017.11.010.

Yoshino, K., Oka, N., Yamamoto, K., Takahashi, H., and Kato, T. (2013). Functional brain imaging using near-infrared spectroscopy during actual driving on an expressway. Frontiers in Human Neuroscience 0, 882. doi:10.3389/FNHUM.2013.00882.

Yücel, M. A., Selb, J., Aasted, C. M., Petkov, M. P., Becerra, L., Borsook, D., et al. (2015). Short separation regression improves statistical significance and better localizes the hemodynamic response obtained by near-infrared spectroscopy for tasks with differing autonomic responses. *https://doi.org/10.1117/1.NPh.2.3.035005* 2, 035005. doi:10.1117/1.NPH.2.3.035005.

Zhang, Y., and Kumada, T. (2018). Automatic detection of mind wandering in a simulated driving task with behavioral measures. PLOS ONE 13, e0207092. doi:10.1371/JOURNAL.PONE.0207092.

Zhu, W., Chen, Q., Xia, L., Beaty, R. E., Yang, W., Tian, F., et al. (2017). Common and distinct brain networks underlying verbal and visual creativity. Human Brain Mapping 38, 2094–2111. doi:10.1002/HBM.23507.

