## Supplementary Table 1 for "Predicting the Degree of Distracted Driving Based on fNIRS Functional Connectivity: A Pilot Study"

Supplementary Material

**Supplementary Table 1:**  Spatial registration of the fNIRS channel location to the AAL atlas.

| **Channel** | **Region** |
| --- | --- |
| 1 | Left precuneus |
| 2 | Right postcentral gyrus |
| 3 | Right postcentral gyrus |
| 4 | Left superior parietal gyrus |
| 5 | Left inferior parietal gyrus |
| 6 | Left middle temporal gyrus |
| 7 | Left postcentral gyrus |
| 8 | Left Paracentral lobule |
| 9 | Left superior frontal gyrus, dorsolateral |
| 10 | Left middle frontal gyrus |
| 11 | Left middle frontal gyrus |
| 12 | Left superior frontal gyrus, dorsolateral |
| 13 | Left superior frontal gyrus, dorsolateral |
| 14 | Left superior frontal gyrus, dorsolateral |
| 15 | Left superior frontal gyrus, medial |
| 16 | Left superior frontal gyrus, medial |
| 17 | Right superior frontal gyrus, medial |
| 18 | Right superior frontal gyrus, dorsolateral |
| 19 | Left middle frontal gyrus |
| 20 | Left superior frontal gyrus, dorsolateral |
| 21 | Left superior frontal gyrus, dorsolateral |
| 22 | Right superior parietal gyrus |
| 23 | Right inferior parietal gyrus |
| 24 | Right middle temporal gyrus |
| 25 | Right percental gyrus |
| 26 | Right postcentral gyrus |
| 27 | Right percental gyrus |
| 28 | Left supplementary motor area |
| 29 | Left supplementary motor area |
| 30 | Right superior frontal gyrus, dorsolateral |
| 31 | Left supplementary motor area |
| 32 | Right middle frontal gyrus |
| 33 | Right middle frontal gyrus |
| 34 | Right superior frontal gyrus, dorsolateral |
| 35 | Right superior frontal gyrus, medial |
| 36 | Right middle frontal gyrus |
| 37 | Right superior frontal gyrus, dorsolateral |
| 38 | Right middle frontal gyrus |
| 39 | Right superior frontal gyrus, dorsolateral |
| 40 | Right middle frontal gyrus |
| 41 | Left superior frontal gyrus, medial |
| 42 | Left superior frontal gyrus, medial |
| 43 | Right superior frontal gyrus, dorsolateral |
| 44 | Right superior frontal gyrus, dorsolateral |
